# High-Throughput Measurement and Machine Learning-Based Prediction of Collision Cross Sections for Drugs and Drug Metabolites

**DOI:** 10.1101/2021.05.13.443945

**Authors:** Dylan H. Ross, Ryan P. Seguin, Allison M. Krinsky, Libin Xu

## Abstract

Drug metabolite identification is a bottleneck of drug metabolism studies. Ion mobility-mass spectrometry (IM-MS) enables the measurement of collision cross section (CCS), a unique physical property related to an ion’s gas-phase size and shape, which can be used to increase the confidence in the identification of unknowns. A current limitation to the application of IM-MS to the identification of drug metabolites is the lack of reference CCS values. In this work, we present the production of a large-scale database of drug and drug metabolite CCS values, assembled using high-throughput *in vitro* drug metabolite generation and a rapid IM-MS analysis with automated data processing. Subsequently, we used this database to train a machine learning-based CCS prediction model, employing a combination of conventional 2D molecular descriptors and novel 3D descriptors. This novel prediction model enables the prediction of different CCS values for different protomers, conformers, and positional isomers for the first time.

---

Drug metabolism studies are a critical component of the drug development process. Metabolites can inform metabolic soft spots and may be pharmacologically active and/or elicit unexpected toxicity or other off-target effects, making knowledge of their structures essential.^1, 2^ Current and conventional approaches to drug metabolite structural determination has typically involved a combination of liquid chromatography (LC), coupled with UV-Vis spectroscopy and/or mass spectrometry (MS), and nuclear magnetic resonance spectroscopy (NMR).^3–5^ LC-MS and LC-UV benefit from low sample requirements and fast analysis time, but identification of unknowns can be limited when relying upon UV spectra or MS fragmentation data alone. In contrast, NMR allows for definitive assignment of chemical structures, but it requires large amounts of materials and is relatively low throughput.

Ion mobility spectrometry (IMS) is an analytical technique that rapidly separates ions based on differences in their gas-phase size and shape, which is orthogonal to polarity-based LC separation and partially orthogonal to mass.^6–10^ In time-dispersive ion mobility (IM) separations, ions are driven through a neutral buffer gas under the influence of an electric field. Ions are differentially impeded as they interact with the buffer gas molecules, and as a result, they traverse the mobility cell in different amounts of time (*i.e.*, drift time). An ion’s drift time can be converted into collision cross section (CCS), a unique physical property reflecting its gas-phase size and shape, using appropriate experimental measurements and/or calibration. Excellent reproducibility has been demonstrated for CCS measured across different instrumentation and labs,^11–14^ making it a robust parameter for compound identification. CCS also provides useful information related to shape, conformation, and polarity. When IM is coupled with MS (IM-MS), an additional dimension of separation is achieved without adversely affecting analytical throughput.

The use of IM-MS for the determination of drugs and their metabolites has gained significant traction in recent years,^15^ but inadequate reference CCS databases remains a significant limitation to the application of CCS to identifying unknown metabolites. Large CCS databases covering drug and drug-like compounds have been presented in the literature,^16–19^ but due to the vastness and complexity of small molecule chemical space, many unknowns may not be represented. This issue of chemical representation is even more pronounced for drug metabolites, for which no such large-scale CCS database exists. This problem can be addressed by leveraging structural trends in existing CCS databases to predict CCS for unknowns that are not in experimental databases, and this approach has been demonstrated by multiple groups, including us.^18–25^ An important consideration in this approach, however, is the dependence of CCS prediction performance on the quality and coverage of chemical space in the data used to train the model.^24^ Therefore, a drug metabolite-specific CCS database is needed for accurate prediction of CCS for drug metabolites. Another limitation in current machine learning (ML)-based CCS prediction models is that the 2D features used in previous work (*e.g.*, molecular quantum numbers, MQNs) do not adequately capture more complex IM behavior arising from the presence of different protomers, conformers, or positional isomers that are common among drugs and drug metabolites.^15^

Here, we present 1) the generation of a high-quality drug and drug metabolite CCS database, through the use of high-throughput *in-vitro* drug metabolite generation and rapid IM-MS analysis with automated data processing, and 2) the training of drug- and drug metabolite-specific CCS prediction models using ML with novel 3D molecular descriptors, which enables the prediction of CCS values for protomers, conformers, and positional isomers with high accuracy and throughput.

## Results

### High-Throughput Measurement of Drug and Drug Metabolite CCS

To obtain a large collection of drug metabolites, we first carried out high-throughput drug metabolism reactions in 384-well plates using human liver microsomes and S9 fraction on 2000 drug and drug-like compounds in the MicroSource Discovery Systems’ Spectrum Collection, containing 50% approved drugs, 30% natural products, and 20% bioactive compounds (Figure 1A). Reactions catalyzed by the Phase-I enzymes, such as cytochromes P450 (CYPs), flavin-containing monooxygenases, and reductases, and Phase-II enzymes, such as glutathione S-transferases (GSTs) and UDP-glucuronosyltransferases (UGTs), were probed. The drug metabolism reactions were carried out with or without enzyme cofactors, such as NADPH (cofactor of CYPs), glutathione (GSH, co-substrate of GSTs), UDP-glucuronic acid (UDPGA, co-substrate of UGTs), and alamethicin^26^ (enabling access of substrates to UGTs). Plates incubated with HLM + S9, but in the absence of enzyme cofactors, served as controls. After the reactions and sample processing, we carried out rapid IM-MS analysis using a 30-mm reverse-phase column, resulting in just under 2 min per run (Figure 1A). Measuring the roughly 2000 compound collection in triplicate with or without enzyme cofactors/co-substrates resulted in > 8900 samples analyzed. This large and complex set of raw data were analyzed using a stepwise approach with a high degree of automation (Figure 1B), including extraction of arrival time distributions (ATDs), gaussian fitting, CCS calibration, and calculation of CCS of observed ATD peaks. The CCS values of parent compounds were obtained by extracting the ATDs of the exact masses of various adducts. For metabolites, we first generated a theoretical list of potential metabolites using Biotransformer,^27^ then extracted the ATDs of these potential metabolites (see Methods for detail). Only ATD peaks meeting the criteria of intensity > 1000 and peak width between 0.06 and 1.77 ms (roughly 1-30 drift time bins) were retained. This approach ultimately led to the assembly of a large CCS database specific to drugs and drug metabolites. Figure 2A summarizes the composition of the drug and metabolite CCS database. The database contained 6245 measured CCS values from 3286 different compounds, of which 1333 were from parent drugs (3675 CCS values) and 1953 were from metabolites (2570 CCS values). The measured CCS values corresponded to a number of ionization states commonly observed in positive mode ESI including [M+H]^+^ (1936), [M+Na]^+^ (1656), [M+K]^+^ (1235), [M+H-H_2_O]^+^ (1299), and [M]^+^ (119).

**Figure 1.**
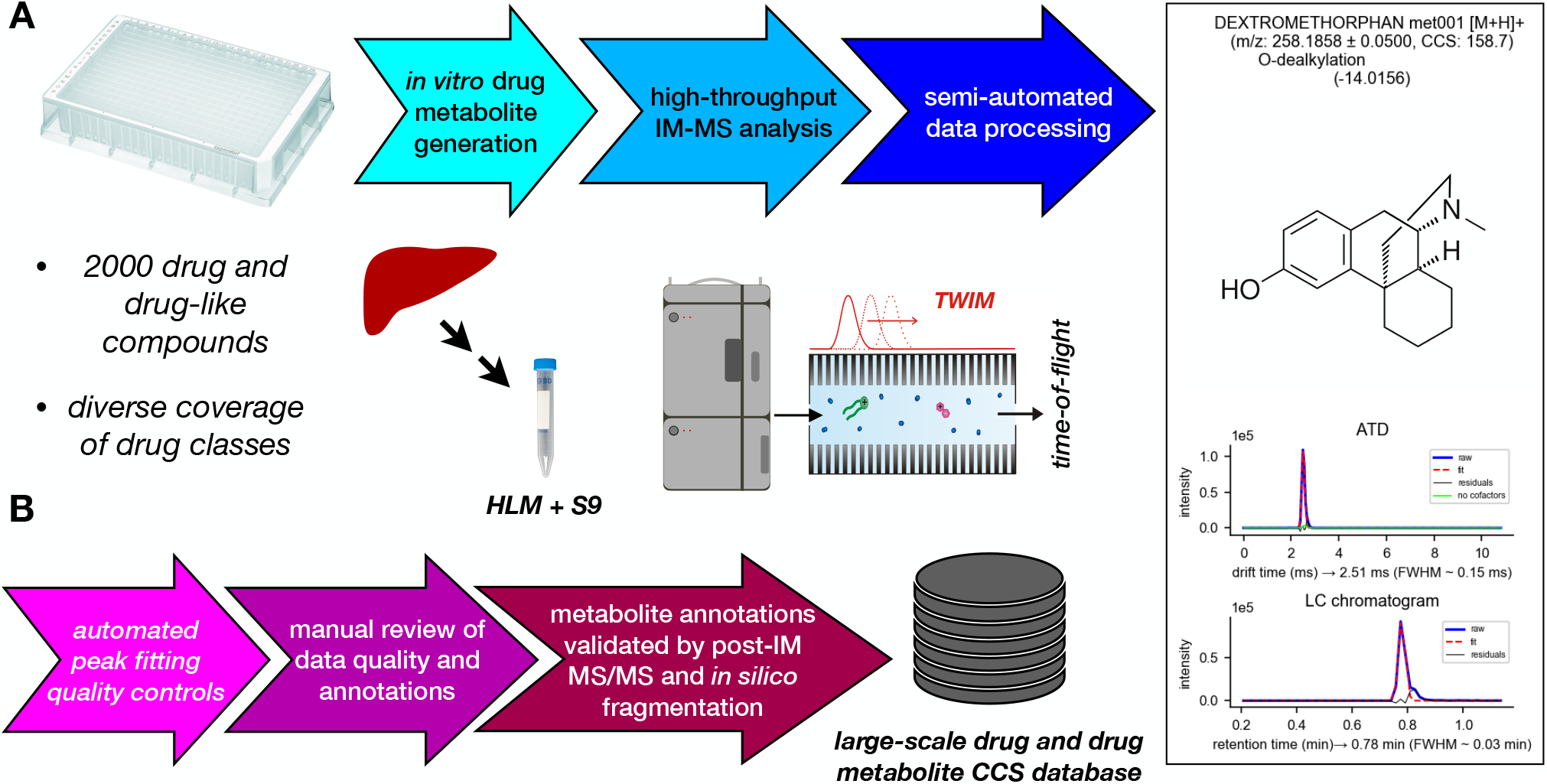
(**A**) Workflow for high-throughput *in vitro* drug metabolite generation and IM-MS analysis. Drug metabolites were generated from the MicroSource Spectrum Discovery Collection, containing ~2000 drug and drug-like compounds, in a high-throughput 384-well plate format using subcellular fractions (microsomes and S9) pooled from 200 human livers. Samples were analyzed using a rapid IM-MS protocol, including semi-automated data processing including extraction and fitting of drift times from ATDs, calibration of CCS, prediction of metabolites, and establishment of cofactor dependence for oxidative metabolites. (**B**) The semi-automated data processing included multiple steps of automated and manual quality controls, including automated quality controls on peak fitting, manual review of extracted data quality and metabolite annotations, and validation of metabolite annotations with MS/MS data. The processed data was finally compiled into a SQLite3 database for use in CCS prediction by ML.

**Figure 2.**
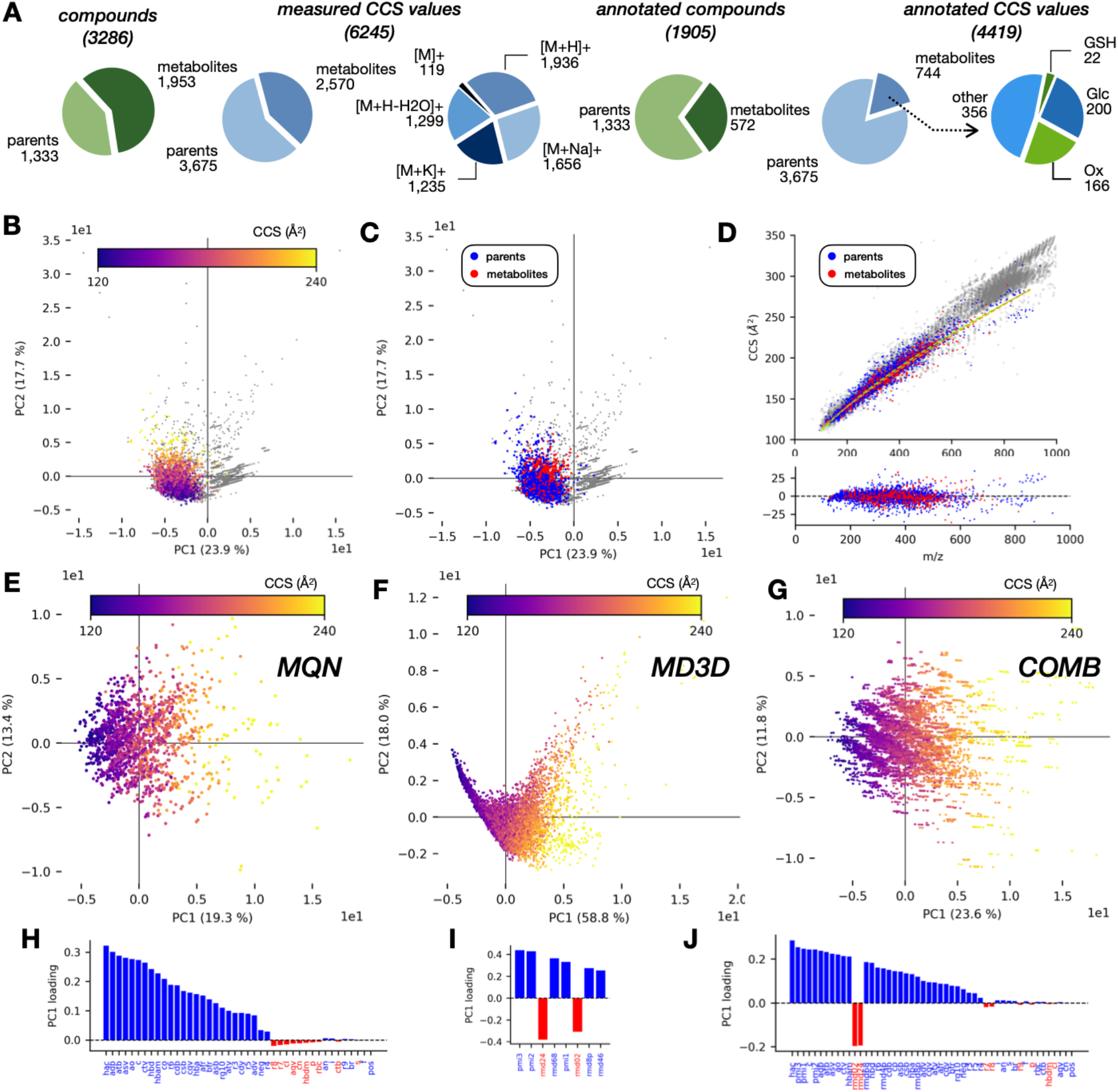
(**A**) Composition of the assembled CCS database for drugs and drug metabolites (dmCCS). Metabolic modification abbreviations: GSH, glutathione conjugated metabolites; Glc, glucuronide metabolites; Ox, oxidative metabolites (*e.g.* −2H, +O, −Me). (**B**) PCA projections of the dmCCS database (color) from a PCA computed using the CCSbase database (grey), colored by CCS. (**C**) PCA projections of parent compounds (blue) and metabolites (red) from the dmCCS database from a PCA computed using the CCSbase database (grey). (**D**) CCS *vs. m/z* of parent compounds (blue) and metabolites (red) from the dmCCS database overlaid on the CCSbase database (grey). Dotted lines represent individual power fits for parent (chartreuse) and metabolite (orange) data, and residual CCS from these fits are included below the main plot. (**E**) PCA projections of dmCCS database computed using MQNs as molecular descriptors, colored by CCS. (**F**) PCA projections of dmCCS database computed using MD3Ds as molecular descriptors, colored by CCS. (**G**) PCA projections of dmCCS database computed using the combination of MQNs and MD3Ds as molecular descriptors, colored by CCS. (**H**) Individual feature loadings for principal component 1 from PCA computed on dmCCS using MQNs as molecular descriptors. (**I**) Individual feature loadings for principal component 1 from PCA computed on dmCCS using MD3Ds as molecular descriptors. (**J**) Individual feature loadings for principal component 1 from PCA computed on dmCCS using the combination of MQNs and MD3Ds as molecular descriptors.

To validate the identity of the potential metabolites, we first carried out a thorough search of DrugBank^28^ for reported metabolites of known drugs in our collection and matched our observed metabolites with those previously reported. For those without reported metabolites, we matched the experimental MS/MS spectra obtained from post-IM fragmentation against an *in-silico* generated MS/MS spectra of potential metabolite structures using MetFrag,^29^ and ruled out low-scoring metabolite annotations. The validation process is discussed in greater detail in Methods. After this process, 4408 of the measured CCS values were retained with an annotation, corresponding to all parent drugs with a CCS value (1333) and 29.3% of metabolites (572). The annotated metabolite CCS values represent a range of metabolic modifications, including glutathione adducts (22), glucuronide conjugates (200), and oxidative metabolites (166). 2D molecular descriptors, *i.e*., molecular quantum numbers (MQNs, 42 features, Table S1),^24, 30^ were generated for all annotated species as described previously. Furthermore, novel 3D molecular descriptors (Table S2), including principal moments of inertia (PMI) and radial mass distributions (RMD) (8 features, see Methods), were generated to better capture the relationship between conformation and CCS during machine learning as discussed below. Briefly, we attempted to generate 3D structures for all annotated [M+H]^+^, [M+Na]^+^, and [M+K]^+^ species at a low level of theory (MMFF94 and PM7, see Methods), resulting in a total of 9813 modeled structures (4074, 3172, and 2567 for each ionized species, respectively). 3D molecular descriptors were generated from 3D structures using in-house developed Python scripts as described in Methods. In total, 7652 and 2161 3D structures with 3D molecular descriptors were generated for parent drugs and metabolites, respectively.

### Characteristics of the Drug and Metabolite CCS Database

Principal components analysis (PCA) was used to probe the chemical characteristics (as captured by 2D or 3D molecular descriptors) that contribute to variance in the drug and metabolite CCS database. We had previously characterized a comprehensive collection of compounds from a diverse set of chemical classes using MQNs as features,^24^ so we first computed a PCA using this comprehensive database (CCSbase) to serve as the chemical space background. We then projected the new drug and metabolite CCS database (dmCCS) into this PCA to examine the chemical space that dmCCS spans within the context of CCSbase. Figure 2B and 2C show the PCA projections of compounds from dmCCS (color) overlaid over compounds from CCSbase (grey). In Figure 2B, it can be seen that CCS values of the compounds from dmCCS generally increase along the direction of PC2, indicating that the strongest sources of variance in CCSbase do not correspond with sources of variance in dmCCS that relate to CCS. Figure 2C shows where the parent compounds and metabolites from dmCCS group fall within the chemical space defined by CCSbase, which indicates that the dmCCS occupies a broad region corresponding roughly to “small molecules”. The metabolites occupy a subspace within the chemical space occupied by the parent compounds. Figure 2D shows where the parent compounds and metabolites from dmCCS (color) map into the IM-MS conformational space (*i.e.* CCS *vs. m/z*), compared to the compounds from CCSbase (grey), with individual power fits for parent compounds and metabolites (dashed lines). Generally, the compounds from dmCCS occupy the low *m/z* region of this space and span a wide range of CCS values. Interestingly, the metabolites seem to occupy a slightly narrower CCS envelope with mostly similar average CCS values to those of the parent compounds. Even in the context of the large chemical space of CCSbase, the compounds from dmCCS represent considerable structural diversity.

Separate PCAs were computed on dmCCS using the 2D (MQN) and 3D (MD3D) molecular descriptors to determine how each set of descriptors reflected the chemical space covered by this database. Figure 2E and 2F show the PCA projections from the 2D and 3D features, respectively. For both feature sets, PC1 correlates well with variation in CCS, indicating that among these compounds the primary sources of variance are related to CCS. PC1 and PC2 of the PCA computed on the 2D feature set captured 19.3% and 13.4% of the overall variance, respectively, compared to 58.8% and 18.0% for the 3D feature set, indicating that a high degree of variance orthogonal to CCS in the 2D feature set is not present in the 3D feature set. Indeed, the PCA computed on the 2D features required 24 components to capture 95% of the variance in the dataset, in contrast to only 5 components needed for the 3D feature set. Figure 2G and 2H show the 2D and 3D feature loadings, respectively, for PC1 in each PCA, both of which correlate well with CCS. The strongest contributors to separation along the first principal component (Figure 2H) for the 2D features were counts of atoms (*hac*: heavy atoms, *ao*: acyclic oxygens, *c*: carbons), bonds (*adb*: acyclic double bonds, *atb*: acyclic triple bonds), and topological features (*asv*: acyclic monovalent nodes, *ctv*: cyclic trivalent nodes). All of the 3D features contributed similarly to the separation along the PC1 (Figure 2I), and interestingly, the second and third PMI had slightly larger contributions than the first. Together, these results demonstrate that both the 2D and 3D features capture the important characteristics of this set of compounds that relate to CCS, but the 3D features contain somewhat less extraneous information.

We next examined the degree to which the 2D and 3D feature sets (MQN and MD3D, respectively) offered complimentary information by computing a PCA on dmCCS using a combination of both feature sets (COMB) (Figure 2G). The PCA projections overall appear quite similar to those from the 2D feature set alone. PC1 captured 23.6% of the total variance, while 27 total components were required to capture 95% of the variance in the dataset. The top features contributing to separation along PC1 (Figure 2J) consist of a combination of those identified from the 2D and 3D feature sets.

We also performed a set of analogous analyses using partial least-squares regression analysis computed on the 2D, 3D, and combined feature sets with CCS as the target variable (Figure S1). The results from these analyses largely mirrored those discussed above, which is expected given the alignment of CCS with PC1 in all three PCAs.

### Training Drug and Drug Metabolite-Specific CCS Prediction Models

The 2D and 3D feature sets were used to train individual ML models for CCS prediction on dmCCS. Despite the different sizes (42 features *vs.* 8) and characteristics of the 2D and 3D feature sets, the MQN and MD3D predictive models achieved very similar performance in CCS prediction by multiple metrics, with robust performance between training and test set data (Figure 3). We next sought to test the degree to which the two feature sets provided orthogonal information by training a ML model on the combined 2D and 3D feature sets (COMB). Although the COMB model achieved significantly improved predictive performance relative to models trained on either feature set (Figure 3), there was a significant lapse in performance between the training and test set data, indicating model overfitting likely attributable to the presence of redundant and/or superfluous features.

**Figure 3.**
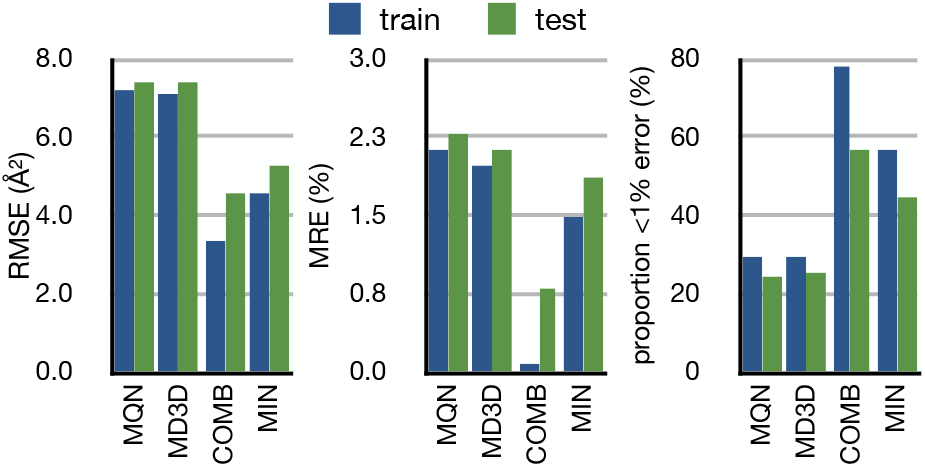
CCS prediction performance comparison for ML models trained on dmCCS using MQN, MD3D, a combination of MQN and MD3D (COMB), or a minimal feature set (MIN) as molecular descriptors.

To address potential overfitting in the COMB model, a set of feature ranking and successive feature removal trials including PLS-RA, gradient boosting regression (GBR), and a permutation feature importance function in *Scikit-Learn* (PER), were run in order to select a minimal feature set combining the most influential features from the 2D and 3D feature sets while avoiding overfitting by removing extraneous features (see Supporting Information for detail). Molecular descriptors retained by at least two of the feature removal methods were kept as the minimal feature set (MIN), which consisted of only 11 descriptors from both the 2D and 3D feature sets: *hac*, *c*, *asv*, *adb*, *ctv*, *hbam* (H-bond acceptor sites), *hbd* (H-bond donor atoms), *pmi1*, *pmi2*, *pmi3*, *rmd02*. A new ML model was trained using this feature set. Although there was still an appreciable degree of correlation between the features (Figure S3), the MIN model achieved an intermediate increase in performance relative to the models trained on the 2D or 3D features alone (Figure 3), and importantly, this performance was better maintained between the training and test set data.

The MIN model was also used to compare with fast theory-based CCS prediction methods (*e.g.* projection approximation, PA, and exact hard-sphere scattering, EHS),^31^ which again showed superior performance in terms of accuracy and precision (see Figure S4 and related text).

### Application of CCS Prediction to Compounds with Multimodal ATDs

Multimodal ATDs can arise from a number of circumstances, such as constitutional isomers (*e.g.*, positional isomers of metabolites or protomers formed in the ESI process) and conformers.^15, 16, 32^ However, previous CCS prediction models based on 2D molecular descriptors generally do not allow the differentiation of such isomers or conformers. Inclusion of 3D features in CCS prediction could in theory capture such multimodal differences for given 3D structures, so we sought to evaluate some known examples of multimodal distributions using CCS prediction models trained with different feature sets.

The oxygenated (+O) metabolite of terfenadine display a more compact conformation relative to the parent, likely attributable to the introduction of an intramolecular polar-polar interaction (Figure 4B).^32^ Figure 4A compares the experimentally measured CCS values of terfenadine and its +O metabolites to values predicted using the different models discussed above. The CCS values predicted using the MQN and MD3D models are lower than the experimental values, and the +O metabolite has a larger CCS than the parent, indicating that these feature sets do not adequately capture the structural differences between these compounds. The MIN model produced the closest predictions, and importantly, reproduced the decreased CCS of the metabolite relative to the parent, with a compaction factor (see Supporting Information) of 1.030 (compared to 1.042 in the experimental values).

**Figure 4.**
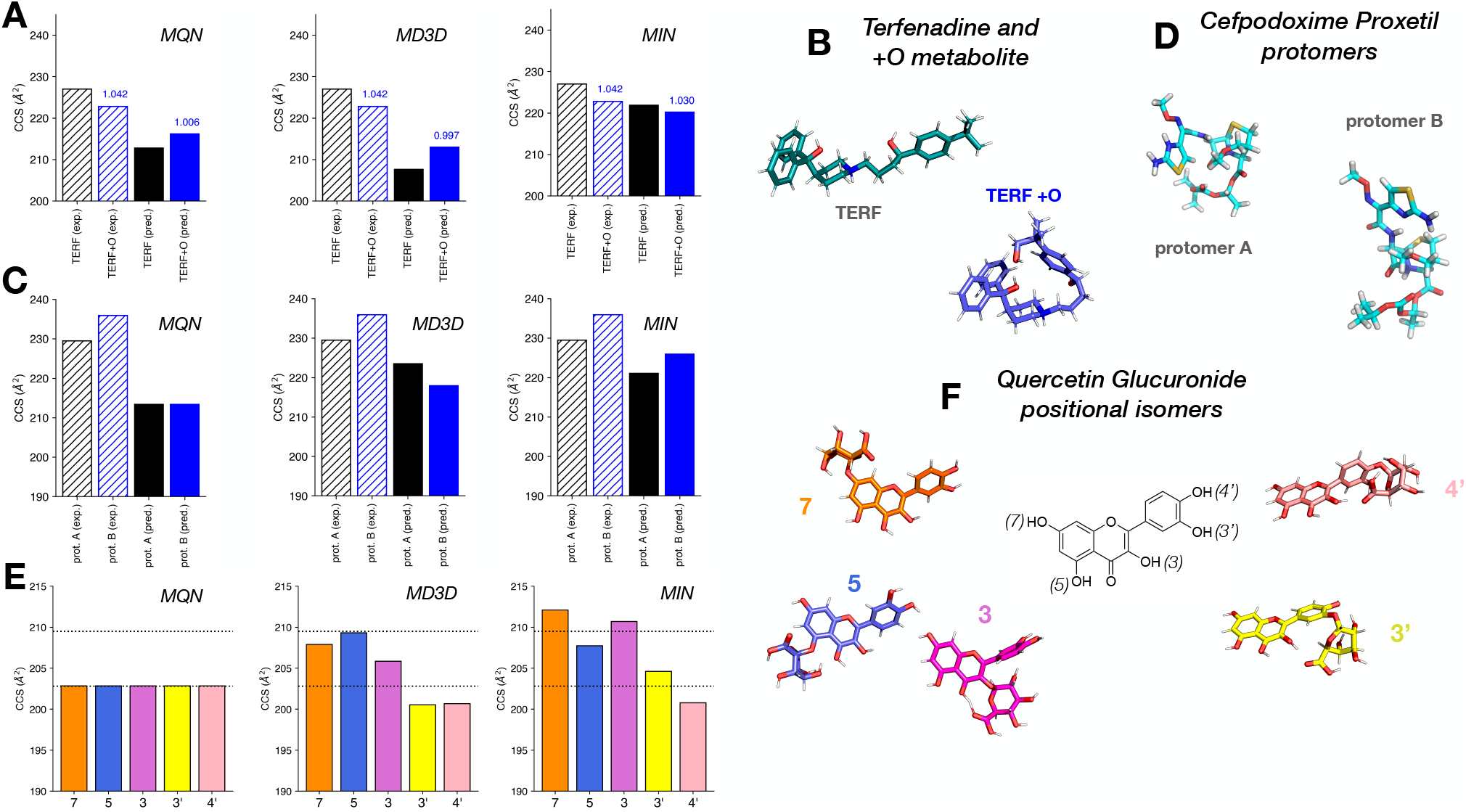
(**A**) Comparison of measured (hatched) and predicted (solid) CCS for terfenadine and its +O metabolite. Each plot presents CCS values predicted using ML models trained on dmCCS using different feature sets. (**B**) Representative structures of terfenadine and its +O metabolite demonstrating the gas-phase compaction of metabolite relative to the parent. (**C**) Comparison of measured (hatched) and predicted (solid) CCS for two protomers of cefpodoxime proxetil. Each plot presents CCS values predicted using ML models trained on dmCCS using different feature sets. (**D**) Representative structures of the two protomers of cefpodoxime proxetil. (**E**) Comparison of measured (dashed lines) and predicted (solid bars) CCS for the positional isomers of quercetin glucuronide. Each plot presents CCS values predicted using ML models trained on dmCCS using different feature sets. (**F**) Representative structures of the positional isomers of quercetin glucuronide.

Cefpodoxime proxetil is a β-lactam antibiotic that has previously been shown to form two protomers in ESI with distinct CCS values (Figure 4D).^16^ As seen in Figure 4C, the MQN model was unable to distinguish between the different protomers (likely due to them being constitutional isomers), and the predicted CCS values were significantly smaller than the experimental values. The MD3D model produced predictions that differed between the two protomers, but their rank-order was reversed relative to the experimental values. The MIN model produced CCS predictions close to the experimental values while preserving the experimentally observed rank-order.

Quercetin is a flavonoid compound with multiple hydroxyl (–OH) groups available for glucuronidation (Figure 4F).^33^ Glucuronidation of quercetin have previously been observed to produce a bimodal CCS distribution, likely attributable to glucuronidation at different positions.^32^ Again, the MQN model failed to capture any CCS differences between the positional isomers while both MD3D and MIN models were able to distinguish between the different positional isomers and the assignment of isomers to the two experimental values was largely in agreement with previous results.^32^

## Discussion

This work addresses several major gaps in applying IM-MS to drug metabolite identification and building ML-based CCS prediction models. First, there is a lack of large-scale experimental CCS database for drug metabolites, which was accomplished here with a high-throughput *in vitro* drug metabolite generation system, followed by high-throughput IM-MS analysis and automated data processing. This large database provided the basis for building ML-based CCS prediction models for drug metabolites. Second, previous ML approaches for the prediction of CCS values rely on 2D molecular descriptors,^18–22, 24^ which cannot differentiate protomers, positional isomers, and conformers. By incorporating novel 3D molecular descriptors, such as PMIs and RMDs, our ML model using minimum combined 2D and 3D features successfully overcame these limitations. Third, our approach represents a hybridization of data- and theory-driven CCS prediction, which showed superior performance than fast theory-based computation approaches in terms of accuracy and precision. Although our ML-based model cannot replace high-level theoretical CCS calculation as small differences captured by the high-level computation methods may not be readily captured by the low-level methods used to generate the training 3D structures in this work, the time-efficiency and accuracy of our approach (a few seconds vs. hours using high-level computation) makes it easily integrated into drug development processes. To summarize, the ML approach reported in this work enables high-accuracy and high-throughput generation of CCS values for drugs and drug metabolites with sufficient precision to differentiate isomers and conformers. The dmCCS prediction models are available to the public at CCSbase.

## Methods

### High-Throughput *in vitro* Drug Metabolite Generation

Drug metabolites were generated *in vitro* using pooled subcellular fractions (S9 and microsomes) derived from human liver following a protocol from our previous work,^32^ adapted to a high-throughput 384-well plate format with all sample preparation performed using automated sample handling systems at the Quellos High-Throughput Screening Core at the University of Washington. Briefly, HLM/S9 stock (5 mM GSH, 5 mM MgCl, 0.01 mg/mL alamethicin, 0.2 mg protein/mL pooled HLM, 0.2 mg protein/mL S9, 100 mM potassium phosphate buffer at pH 7.4) was prepared and allowed to stand on ice for 15 min (alamethicin pre-treatment to enhance UGT activity). 90 μL of the HLM/S9 stock was dispensed into each well of 14 384-well plates, then 0.5 μL of each drug stock (50 mM in DMSO) from the MicroSource Spectrum Discovery Collection (7 plates) were dispensed into the plates in duplicate. 10 μL of a cofactor-containing activation mixture (10 mM NADPH, 50 mM UDPGA, 100 mM potassium phosphate buffer at pH 7.4) or potassium phosphate buffer without cofactors (as control) were then added to the duplicate plates, initiating the drug metabolism reactions for plates containing activation mixture. All plates were incubated at room temperature for 90 min before being quenched with 100 μL ice-cold acetonitrile (with 10 μM lysophosphatidylethanolamine 13:0 as an internal standard). After quenching, all plates were stored at 4 °C for at least 15 min to promote precipitation of proteins. Each plate was centrifuged at 3500G for 15 min at 4 °C to sediment the precipitated proteins, then 150 μL of the supernatant was transferred to fresh plates. All plates were stored at −80 °C until IM-MS analysis.

### High-Throughput Ion Mobility-Mass Spectrometry

Samples (5 μL) were injected and separated using a Waters Acquity FTN UPLC coupled to a reverse-phase column (Phenomenex Kinetex, 2.6 μm, polar C_18_, 100 Å, 30 × 21 mm), eluting with a gradient of water with 0.1% formic acid (A) and methanol with 0.1% formic acid (B) at 0.5 mL/min: 0.00-0.20 min, 100% A; 0.20-0.30 min, 100→25% A; 0.30-0.75 min, 25→0% A; 0.75-1.05 min, 0% A; 1.05-1.10 min, 0→100% A. The total analysis time for each sample, factoring in acquisition and autosampler operations, was just under 2 min. For each injection, the first 0.20 minutes of eluent was diverted to waste in order to avoid buildup of salt on the ESI source, and after that, the flow was automatically diverted back to the instrument via an electronically controlled switching valve. TWIM-MS analysis was performed on a Waters Synapt G2-Si mass spectrometer (Waters Corp., Milford, MA) equipped with an ESI source and using nitrogen as the drift gas. ESI conditions were as follows: capillary, +2.3 kV; sampling cone, 40 V; source temperature, 130 °C; desolvation temperature, 350 °C; cone gas, 90 L/h; and desolvation gas, 600 L/h.

Mass calibration was performed using sodium formate for the range of *m/z* 50−1200. IM separations were performed at a traveling wave velocity of 650 m/s and a height of 24.9 V. For post-IM fragmentation analyses, collision energy was added to the transfer region using a ramp from 30 to 50 eV. Data was acquired from 0.20 to 1.15 min with a 1s scan time over m/z 50−1200, which resulted in approximately 57 scans across the acquired elution region (individual peaks typically spanned ~0.05 min for roughly 3 scans per peak). The 384-well plates were analyzed on three separate occasions over two months.

### TWIM CCS Calibration

A series of singly charged polyalanines (n = 2−14) and a mixture of drug-like compounds were used for calibration of TWIM drift times into CCS (^TWIM^CCS_N2_) using their drift tube CCS values in nitrogen (^DT^CCS_N2_), as described previously.^16, 32^ Briefly, arrival time distributions (ATDs) for CCS calibrants were extracted from the raw data (acquired multiple times throughout acquisition of each plate) using accurate mass with a window of ±0.01 Da, and a CCS calibration curve was constructed from reference CCS values in an automated fashion using a Python script developed in-house.^32^ Drift times for each calibrant were obtained as the mean from a least-squares fit of a Gaussian function on the ATD and were corrected for mass-dependent flight time outside the mobility region to give the corrected drift times (t_d_′), and reference CCS values were corrected for the ion charge state (Z) and reduced mass with the drift gas to give the corrected CCS (CCS′).^34^ A calibration curve was generated by fitting these corrected values with the function CCS′ = A(t_d_′ + t_0_)^B^, where A, t_0_, and B were the fitted parameters.^11, 35^ A calibration curve displaying randomly distributed fit residuals with a maximal absolute error of less than 3% was considered acceptable. CCS calibrant data was acquired 3-5 times over the course of analysis of each plate, and all of this calibrant data was used to construct a combined CCS calibration curve to account for any variation that occurred over the course of data acquisition.

### Ion Mobility-Mass Spectrometry Data Processing

The raw IM-MS data was processed in a number of steps to extract, annotate, and validate CCS values for drugs and putative metabolites (Figure 1B), and was performed separately for each batch of data acquired on the same day (two plates were analyzed each day). The first set of data processing steps were completely automated using Python scripts developed in-house and were applied only to the first technical replicates. First, a target list was assembled for the parent drugs based on the known plate contents. For each parent compound, *m/z*-selected arrival time distributions (ATDs) were extracted for common ionized species with a tolerance of 0.05 Da. ATDs were fit with a gaussian function to obtain drift time, and the fitted drift time was used to calculate calibrated CCS. If an ATD was not able to be fit or the fitted peak did not meet empirically determined rough quality cutoffs (intensity > 1000, peak width between 0.06 and 1.77 ms), the corresponding ionized species was not processed any further. Upon successful ATD peak fitting, a drift time-selected chromatogram was also extracted and an attempt was made to fit for retention time. All data and metadata were stored in custom Python data structures for subsequent processing. Putative metabolites were generated using BioTransformer^27^, with the “allHuman” setting and up to 2 metabolism steps. Putative metabolites were filtered to exclude isobaric metabolites, metabolites with the same neutral mass as the parent compound, and metabolites resulting from the breakdown of secondary metabolites (*i.e.* free glucuronic acid or glutathione), then their corresponding ATDs were extracted from the raw data and fitted as described above. Successfully fitted ATDs were stored along with metadata (including putative metabolite annotation) in custom Python data structures for subsequent processing. Plots containing compound/putative metabolite structures, *m/z*, metabolism reaction information, CCS, and ATDs with fits were generated and stored (see Figure 1A) for subsequent manual review.

The resulting initial data set (>11k analytes) was next subjected to a manual review process. Each of the generated plots described above were manually inspected for general quality of ATD peak fitting (clean ATD fit without secondary peaks) and cofactor-dependence for oxidative and glucuronide metabolites, then accepted or rejected accordingly. The results of this manual review process were used to curate an analyte *m/z* target list for automated data extraction from the second and third replicates. Data extraction from the second and third replicates followed the same automated workflow described above, except that the curated target list was used to search for putative metabolites rather than through *in silico* metabolite prediction. All extracted data and metadata from the second and third replicates were stored in custom Python data structures for subsequent processing.

The final step in data processing was validating compound annotations, which was performed using a semi-automated process. The identities of the parent compounds were known from the plate contents, so further validation was not required. To validate the annotations of the putative metabolites, known metabolites of the parent compounds were manually searched for in the DrugBank database.^28^ A list of potential metabolites and associated metadata were compiled from these searches and later matched to metabolites (superseding the original putative metabolite annotation from BioTransformer) on the basis of their neutral mass (within 50 ppm was considered a match). Finally, all metabolite annotations were subjected to filtering based on post-mobility MS/MS data that were acquired for the first replicate. Drift time-selected MS/MS spectra were extracted and scored against *in silico* fragmentation spectra using MetFrag,^29^ and all annotations with a fragmenter score above the empirical cutoff of 100 were accepted. This empirical MetFrag scoring cutoff was determined by a rank test using the known identities of the parent compounds as follows. The drift time-selected MS/MS spectrum for each parent compound was compared to the *in silico* fragmentation spectra of all parent compounds, resulting in ranked identifications with corresponding fragmenter scores. The rank and score of the known identity were recorded for each compound, and the empirical scoring cutoff was determined by looking at the distribution of scores for parent compounds with true identities ranking in the top 500 (Figure S5A). Ultimately, this cutoff represents a rough way of ruling out unlikely annotations given their corresponding MS/MS spectrum, and in total 861 putative metabolite annotations were removed based on this criterion (587 annotations without scores + 274 with scores < 100).

### Assembly of a Drug and Metabolite CCS Database

A SQLite3 database was used to store experimental data, associated metadata, annotations, 3D structures, and computed molecular descriptors for all of the drugs and metabolites observed in this study. The overall database architecture is summarized in Figure S6. The database has separate tables for CCS measurement data and metadata (*plate_N*), MS2 spectra (*plate_N_ms2*), compound annotations (*plate_N_id*), 2D molecular descriptors (*plate_N_mqn*), 3D structures (*plate_N_3d*), and 3D molecular descriptors (*plate_N_md3d*). All of the experimental plates (7 in total) have their own set of corresponding tables for consistency with the organization of the experimental source data. The database was constructed in a stepwise, automated fashion using a series of Python build scripts developed in-house. Briefly, the database was first initialized with all of the empty tables, then the measured data were added to the *plate_N* tables according to plate number. Next, compound annotations (names and SMILES structures) were added to the *plate_N_id* tables. Parent drug annotations were already known from the plate contents, but metabolite annotations were assigned via a combination of automated and manual processes as discussed above. Drift time-selected MS^2^ spectra were added to the *plate_N_ms2* tables, with each entry in the measured data tables having a corresponding MS^2^ spectrum. Next, MQNs were computed using SMILES structures (see below for detail) from the annotation tables and added to the *plate_N_mqn* tables. 3D structures (in plain text format) were generated (see below for detail) and added to the *plate_N_3d* tables, and corresponding 3D molecular descriptors were computed for each structure (*vide infra*) and added to the *plate_N_md3d* tables. The *plate_N*, *plate_N_ms2*, and *plate_N_id* tables are all related by a unique (across all 7 sets of plates) text identifier, *dmim_id*. All of the annotations in the *plate_N_id* tables have an additional unique integer identifier, *ann_id*, relating them to entries in the *plate_N_mqn* and plate_N_3d tables. The *plate_N_3d* tables have an additional unique integer identifier, *str_id*, relating their entries to the *plate_N_md3d* tables.

### Generation of 3-Dimensional Structures for Ionized Drugs and Metabolites

3-Dimensional structures were computed from SMILES structures for experimentally observed ionized (protonated and Na^+^/K^+^ adducts) drug and metabolite species using a series of scripts developed in-house employing a combination of molecular mechanics and semi-empirical methods. Briefly, initial 3D structures were generated by a Monte Carlo conformer search followed by steepest descent energy minimization using the MMFF94 force field in the OpenBabel^36^ software package. The initial 3D structures were then further optimized at the PM7 semi-empirical theory level in Gaussian16^37^. Finally, the optimized atom positions, masses, and partial charges were stored along with relevant metadata for the measured species. This process was repeated 3 times for each individual ion species to increase the chances that a minimum energy structure would be sampled in this non-extensive modeling protocol.

Generation of 3D structures for protonated species followed the same protocol, but with the inclusion of additional steps to account for multiple potential sites of protonation within a molecule. First, potential protomers were determined by presence of ionizable groups, and the SMILES structures were modified to reflect each protomer. 3D structure generation was performed using each of the protomer SMILES structures as described above, but with additional thermodynamic calculations specified in the semi-empirical optimization step. After 3D structures had been produced for all potential protomers, the structures having the lowest energy and highest partial charge located on the protonation site (if different from lowest energy structure) were selected and stored. The above protocol was repeated 3 times for each species, resulting in 3-6 structures for each protonated species.

The 3D structure generation protocol resulted in the production of 3-6 structures for each ionized species in an attempt to capture multiple energetically similar conformers; however, for most compounds, many or all of the produced structures were virtually the same. To avoid undue influence in predictive model training from such duplications, all structures for a given compound were subjected to RMSD filtering. Briefly, for each compound, a mass-weighted RMSD matrix was computed between all predicted structures and only those differing by more than 0.01 Å were retained. The RMSD cutoff of 0.01 Å was determined empirically by computing the distribution of RMSD values for all structures in the database (Figure S7A), in addition to manual inspection of a handful of compound structures. All of the filtered 3D structures were added as a separate table to the drug and metabolite CCS database.

### Generation of 2-Dimensional Molecular Descriptors

Molecular quantum numbers^*38*^ (MQNs) were used as 2D molecular descriptors for analysis of the drug and metabolite CCS database. MQNs are graph properties of a 2D molecular structure (e.g. a SMILES structure), which include counts of atoms, bonds, and topological features. MQNs were computed from the neutral SMILES structures for all entries in the drug and metabolite CCS database using the RDKit library (https://www.rdkit.org). The computed MQNs were added as a separate table to the drug and metabolite CCS database.

### Generation of 3-Dimensional Molecular Descriptors

Principal moments of inertia (PMI) and binned radial mass distributions (RMD) were used as molecular descriptors for 3D molecular structures. PMI are derived from the eigendecomposition of the inertia tensor of a rigid body computed relative to its center of mass. Physically, this computation produces a set of orthogonal axes within a body, such that the radial distribution of mass about each successive axis is minimized; the magnitude of the PMIs reflects the extent of radial mass distribution about their corresponding axes (Figure S7B). Given a 3D molecular structure defined by N atoms having masses (m) and positions (x, y, z) with center of mass located at the origin, the body frame inertia tensor (I) was computed as follows:

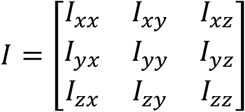

where the diagonal elements (*I_xx_*, *I_yy_*, *I_zz_*) were computed as:

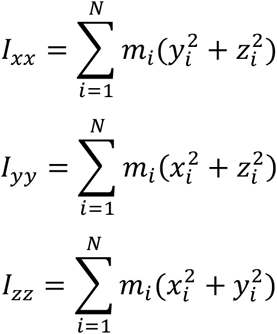

and the off-diagonal elements (*I_xy_*, *I_yx_*, *I_xz_*, *I_zx_*, *I_yz_*, *I_zy_*) were computed as:

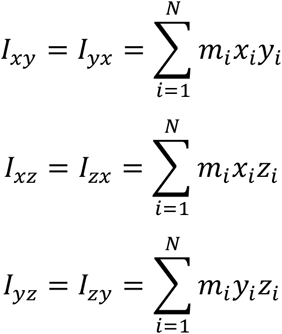

An eigendecomposition (as implemented in the SciPy^39^ Python library: *scipy.linalg.eigh*) was then performed on the inertia tensor, yielding the principal moments of inertia (*PMI_1_*, *PMI_2_*, *PMI_3_*):

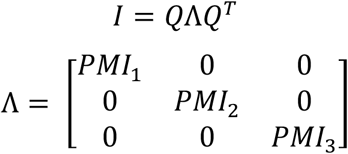

RMDs reflect the proportions of a structure’s mass that lie within specific distances radially from its center of mass. Specifically, RMDs are normalized, mass-weighted histograms of atomic distances relative to the center of mass. The histograms were binned at specific distance intervals (0-2 Å, 2-4 Å, 4-6 Å, 6-8 Å, and >8 Å) in order to reduce the total number of features. The binning intervals were chosen based on the combined distribution of mass-weighted radial distances from all 3D structures in the drug and metabolite CCS database (Figure S7C). The computed 3D molecular descriptors were added as a separate table to the drug and metabolite CCS database.

### Multivariate Analysis of Drug and Metabolite CCS Database

PCA and PLS-RA are implemented in *Scikit-Learn*, a free and open-source machine learning library for Python (*sklearn.decomposition.PCA* and *sklearn.cross_decomposition.PLSRegression*, respectively)^40^. PCA and PLS-RA are dimensionality reduction techniques that work by determining successive orthogonal axes within a high-dimensional dataset that contain maximal variance. PLS-RA differs from PCA in that the first axis is chosen such that it corresponds to the direction of maximal variance in an external target variable (in this case CCS), making it a targeted analysis.

### Feature Selection for CCS Prediction

Starting from a complete combined feature set (2D + 3D molecular descriptors, 50 features total), a set of tests were performed (using only the training set data) to determine the minimal feature set necessary to make robust and accurate CCS predictions. First, the relative importance of all individual features was determined by three methods: PLS-RA, gradient boosting regression (GBR, *sklearn.ensemble.GradientBoostingRegressor*), and a permutation feature importance function built into *Scikit-Learn* (PER, *sklearn.inspection.permutation_importance*). PLS-RA gives an indication of feature importance based on the magnitude of the loadings in the x-dimension (*i.e.* the multidimensional axis that explains the maximal variance in the target variable). GBR is an ensemble method in which successive decision tree models are fitted to the residuals of previous models, and relative feature importance can be inferred from the frequency with which individual features are used for decision tree splits. In the PER method, feature importance is related to the decrease in prediction performance when a feature is randomly shuffled relative to a baseline (unshuffled) performance. Once feature importance had been calculated, sequential feature removal tests were performed using the importance from each method. In the feature removal tests, the least important features were successively removed, and new predictive models were trained and evaluated on the smaller feature sets. This process was repeated until only a single feature (with the highest importance) remained (Figure S2A-C). For each method, a reduced feature set was selected as the set of features for which the prediction error (RMSE) increased above 5 Å^2^ upon their removal. Finally, a minimal feature set was selected as those common among 2 sets of features remaining after feature removal tests using the PLS-RA, GBR, and PER feature importance (Figure S2D).

### Prediction of CCS Using Machine Learning

Prior to model training, the data were processed in a stepwise fashion. First, the data set was randomly (seeded for deterministic results) split into training and test sets in proportions of 80% and 20%, respectively, and the test set was held aside during model training. Rough stratification based on distribution of CCS was used during data set splitting to ensure comparability between the training and test sets. The training data were centered and scaled such that each feature would have a mean of 0 and unit variance in order to avoid undue emphasis of features on the basis of their magnitudes. A support vector regression (SVR) model with radial basis function kernel was used for CCS prediction (*sklearn.svm.SVR*). The model hyperparameters (*C* and *gamma*) were optimized using a grid search with 5-fold cross validation (*sklearn.optimize.GridSearchCV*) on the training data. The model trained using the optimal hyperparameters was then used to compute performance metrics (*vide infra*) from predictions made on the training and test data sets.

### CCS Prediction Performance Metrics

A standard set of metrics were used to determine the bulk performance of CCS prediction using ML and other methods as described previously^24^. Briefly, these include R^2^, mean and median absolute error (MAE and MDAE, respectively, Å^2^), mean and median relative error (MRE and MDRE, respectively, %), root mean squared error (RMSE, Å^2^), and cumulative error distribution at 1, 3, 5 and 10% levels (CE135A, %).

### Calculation of PA/EHS CCS

Theoretical CCS values were calculated for all 3D structures in the drug and metabolite CCS database by the projection approximation (PA) and exact hard-sphere scattering (EHS) methods using MobCal.^31, 41^

### Calculation of Metabolite Compaction Factors

Gas-phase compaction factors of metabolites relative to parents were computed using the equation:

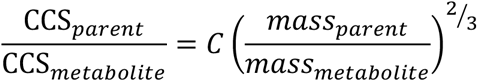

as described previously.^32^ Briefly, since the CCS at a given mass is analogous to a gas-phase density, a change in mass (*i.e.* due to metabolic modification) is expected to produce a monotonic change in CCS, and that change is isotropic if the density does not change (C = 1). If C > 1, then the metabolite is denser than expected under isotropic growth while C < 1 indicates the metabolite is less dense.

## Supporting information

Supplementary Inforamtion

## Data Availability Statement

The prediction models are available at CCSbase.net. Raw data is available upon request from the authors.

## Code Availability Statement

All codes for generating the database and prediction models and for data processing are available at Github (https://github.com/dylanhross/dmccs).

## Acknowledgements

This work was supported by the University of Washington (UW) CoMotion Innovation Gap Fund Chemistry to L.X. We thank Prof. Matthew F. Bush at UW Chemistry for helpful discussion on CCS calculation using MobCal.

## Author Contributions

D.H.R. and L.X. designed the study; D.H.R. and R.P.S. developed *in vitro* drug metabolite generation methodology; D.H.R. developed and performed IM-MS analysis; D.H.R. designed and conducted semi-automated data analysis; D.H.R. performed ML experiments with contributions from A.K.; D.H.R. and L.X. prepared the manuscript with contributions from R.P.S.; All authors approved the submitted version of this manuscript.

