## Supplementary Inforamtion for "High-Throughput Measurement and Machine Learning-Based Prediction of Collision Cross Sections for Drugs and Drug Metabolites"

#### 1. Supplementary Results

*1.1 – Partial Least-Squares Regression Analysis on dmCCS*

Figure S1 shows the results from partial least-squares regression analyses (PLS-RA) computed on dmCCS using 2D, 3D and combined feature sets with CCS as the target variable. Figures S1A-C show the PLS-RA projections of the dmCCS database computed using 2D, 3D, and combined molecular descriptors, respectively. When compared against the corresponding PCA projections (Figures 2E-G in the main text), the overall distributions are similar. Further, the x-loadings from the PLS-RAs (Figures S1D-E) display nearly identical rank-ordering relative to the PC1 feature loadings (Figures 2H-J in the main text). Taken together, these results show that a targeted analysis of dmCCS reproduces the same basic conclusions as those garnered from PCA for all feature sets tested, which is to be expected given the high degree of alignment between CCS and PC1 observed in the PCAs.

*1.2 – Comparison of CCS Prediction Model to Theory-Based Conventional Methods*

Computational modeling to produce 3D structures at a low theory level is the primary bottleneck in training and application of ML models for CCS prediction based on 3D molecular descriptors. Given that production of such structures is also a bottleneck for some of the faster theory-based CCS prediction methods (*e.g.* projection approximation, PA, and exact hard-sphere scattering, EHS),[1] we sought to compare the accuracy of CCS values predicted using both approaches for compounds in dmCCS. Figure S4A shows measured and calculated CCS for compounds from dmCCS, colored according to calculation method. The ML values were predicted using the model trained on the MIN feature set described in the previous section. Both of the theory-driven methods (PA and EHS) display significant systematic errors; however, these systematic errors are likely attributable to the parameterization of these methods, which were originally optimized for He as the drift gas. When systematic errors were corrected using linear regression, the residuals of the fit for PA or EHS-generated values were significantly larger than the ML-predicted values (Figure S4). Taken together, it is clear that ML-based CCS prediction produces higher quality CCS values with this dataset than comparable theory-based methods, likely attributable to the nuanced structural trends that such ML model can capture when provided with appropriate training data.

### 2. Supplementary Figures and Tables

#### *2.1 – Figure S1: Partial Least-Squares Regression Analysis on dmCCS*

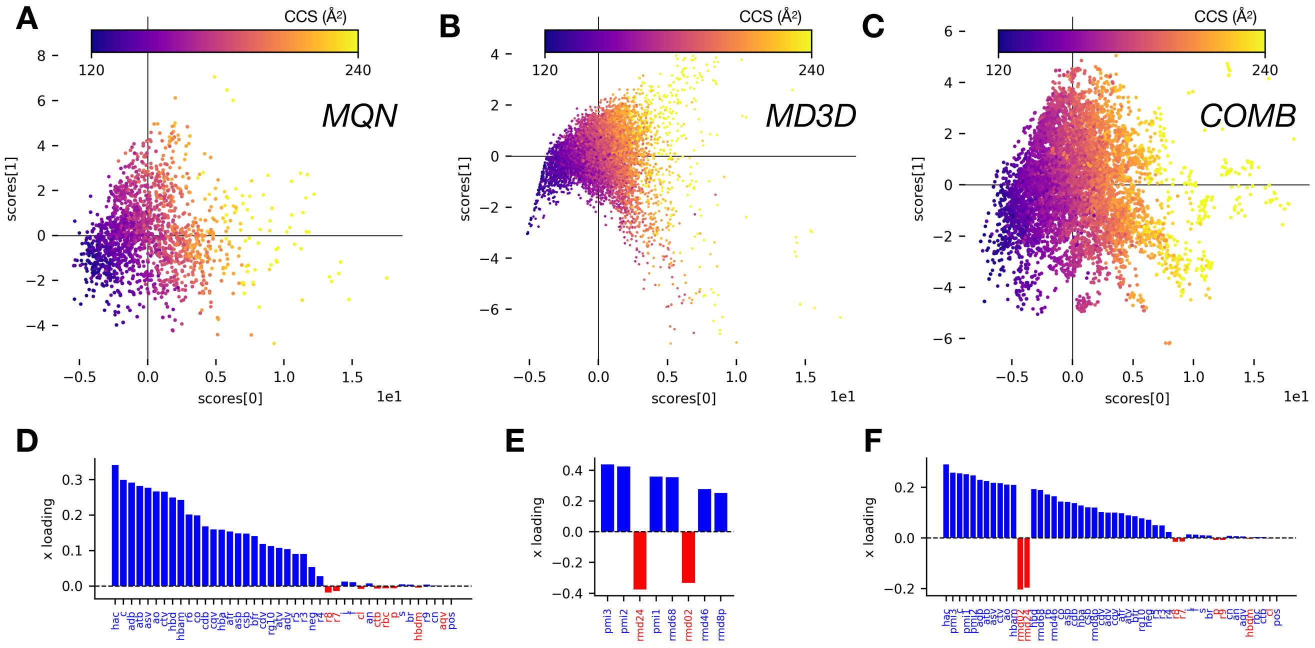

**Figure S1.** (**A**) PLS-RA projections of dmCCS database computed using MQNs as molecular descriptors and CCS as the target variable, colored by CCS. (**B**) PLS-RA projections of dmCCS database computed using MD3Ds as molecular descriptors and CCS as the target variable, colored by CCS. (**C**) PLS-RA projections of dmCCS database computed using the combination of MQNs and MD3Ds as molecular descriptors and CCS as the target variable, colored by CCS. (**D**) Individual feature loadings for component 1 from PLS-RA computed on dmCCS using MQNs as molecular descriptors and CCS as the target variable. (**E**) Individual feature loadings for component 1 from PLS-RA computed on dmCCS using MD3Ds as molecular descriptors and CCS as the target variable. (**F**) Individual feature loadings for component 1 from PLS-RA computed on dmCCS using the combination of MQNs and MD3Ds as molecular descriptors and CCS as the target variable.

*2.2 – Figure S2: Feature Selection Trials*

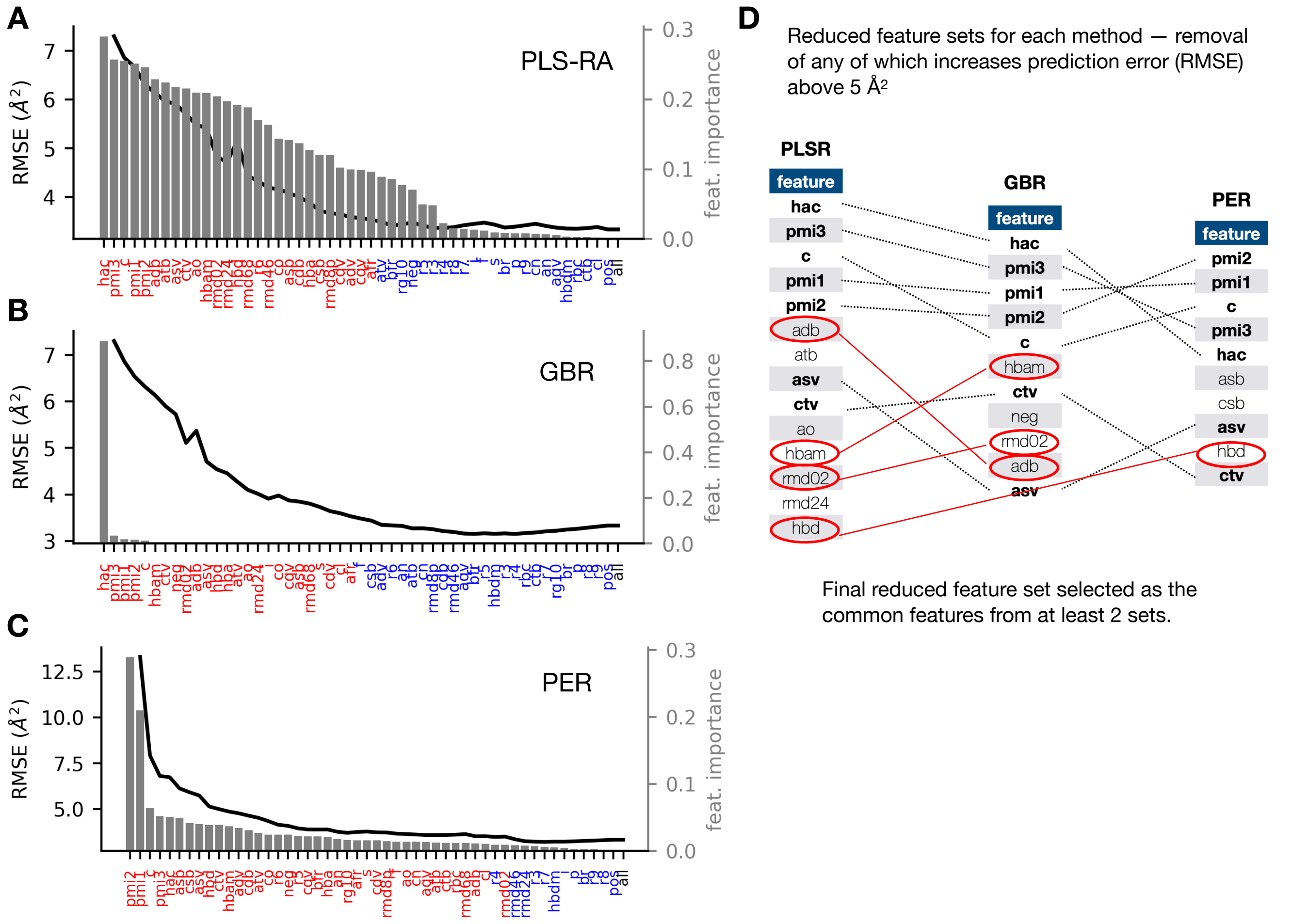

**Figure S2.** (**A**-**C**) Results from feature selection trials. Features were removed in descending order of feature importance (from right to left, grey bars), and resulting predictive model performance was recorded (RMSE, black line). Blue labels indicate the features that could be removed without model performance increasing RMSE more than 5% relative to the baseline (*all*). (**D**) Selected features from individual trials, selected as those for which removal increased error above 5 Å². The features selected in at least two of the individual tests were retained as the final minimal feature set.

*2.3 – Figure S3: Minimal Feature Set Correlation Matrix*

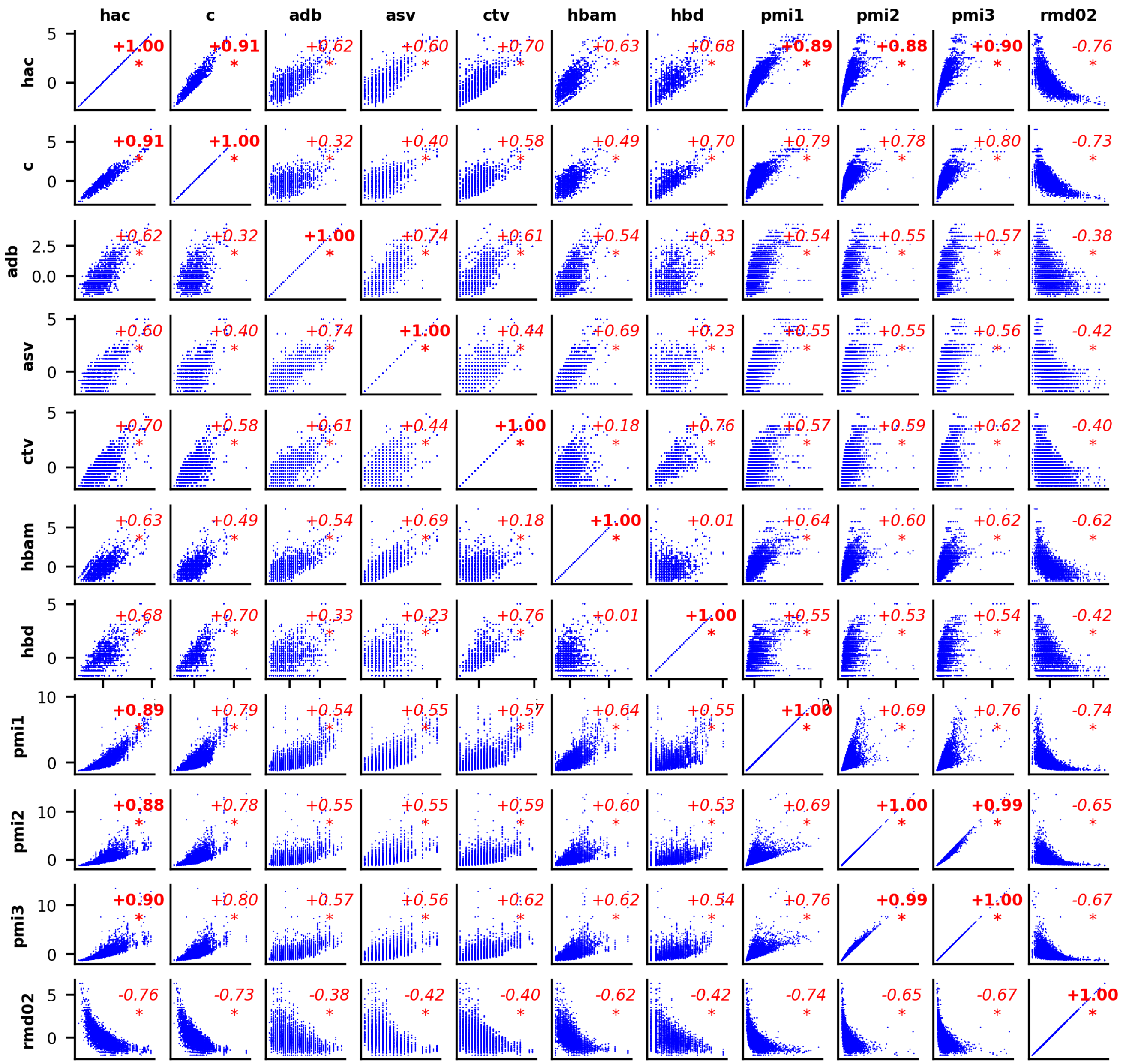

**Figure S3.** Correlation matrix of minimal feature set from feature selection trials. Red numbers correspond to spearman rank test correlation coefficients (coefficients with magnitude > 0.85 are in bold). Asterisks denote a p-value < 0.01 for the correlation.

*2.4 – Figure S4: Comparison of measured CCS and CCS predicted using PA/EHS methods*

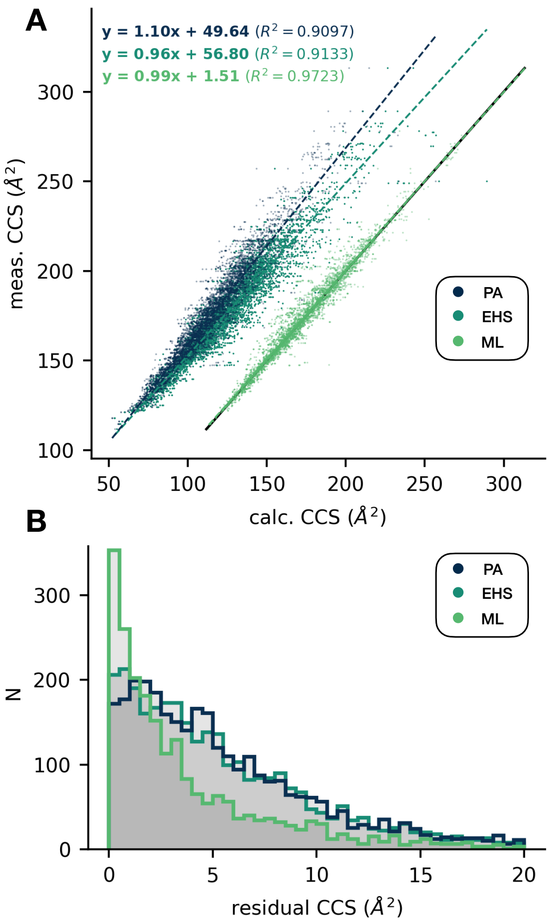

**Figure S4.** (**A**) Comparison of measured CCS and CCS predicted using PA/EHS methods or by a ML model trained on the dmCCS database. Dotted lines correspond to linear fits on each set of values. (**B**) Distributions of residuals from each linear fit.

#### *2.5 – Figure S5: MetFrag Fragmenter Score Cutoff*

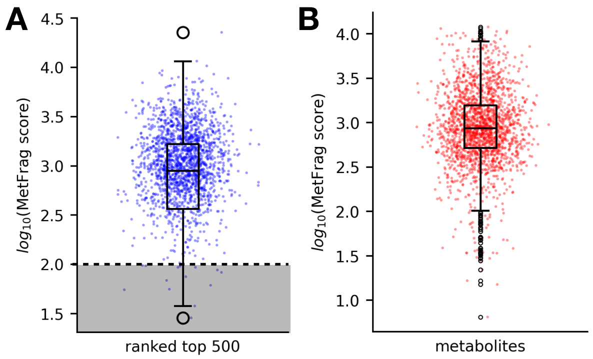

**Figure S5.** (**A**) Distribution of log-transformed MetFrag fragmenter scores for all parent compounds with true annotations ranked in the top 500 from the parent rank test. The dashed line indicates the empirically determined cutoff used to filter out metabolite annotations during construction of the dmCCS database. (**B**) Distribution of log-transformed MetFrag fragmenter scores for metabolites in dmCCS prior to filtering. The center line is the median, the box edges are the upper/lower quartiles (*i.e.*, Q1 and Q3), the whiskers are 1.5x the interquartile range, and the points are outliers beyond the whiskers.

#### *2.6 – Figure S6: dmCCS Database Architecture and Composition*

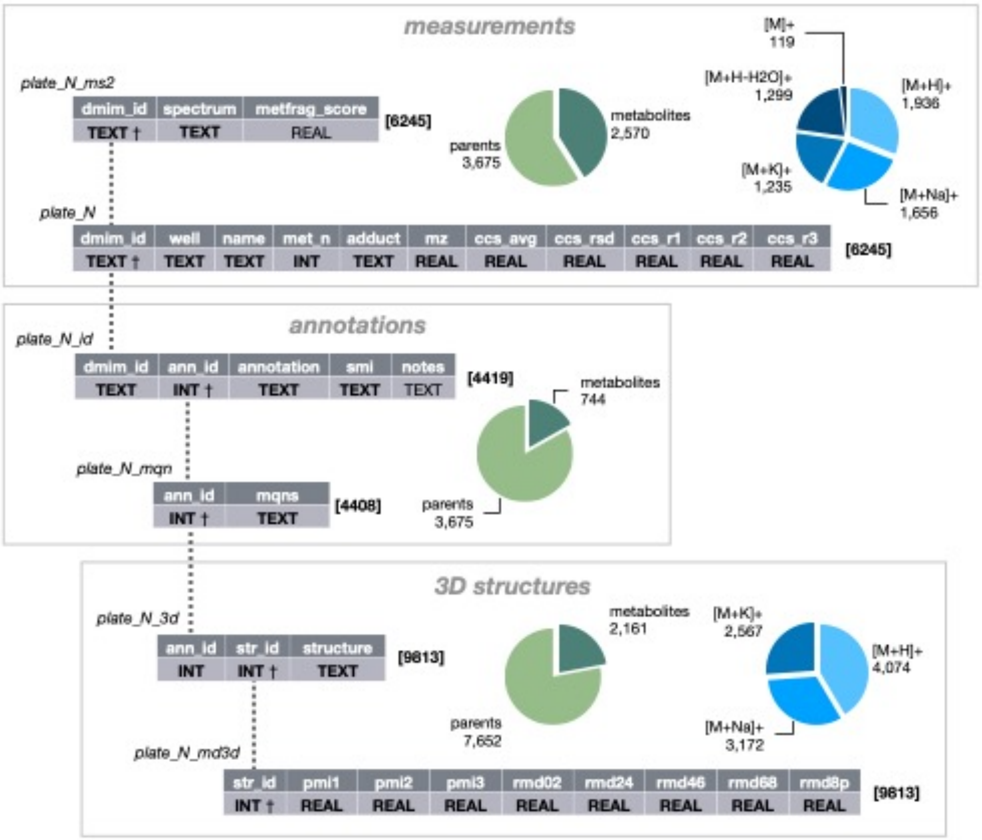

**Figure S6.** Overview of the structure of the dmCCS SQLite3 database. Each grey box represents the general type of information contained within each table, and pie charts reflect characteristics of these grouped tables.

The names and data types are shown for each table, with bold datatypes indicating a required column and † indicating the primary key of the table. The dashed lines indicate the related columns between each table.

##### *2.7 – Figure S7: Description of 3D Molecular Descriptors*

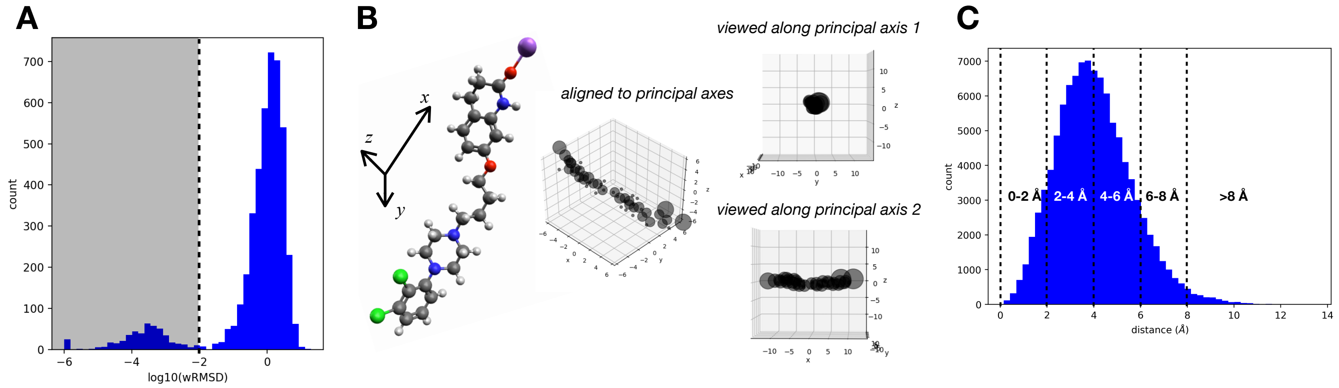

**Figure S7.** (**A**) Distribution of log-transformed mass-weighted RMSD for all pairwise combinations of multiple 3D structures for all compounds in the dmCCS database. The dashed line indicates an empirically determined cutoff used for determination of whether individual 3D structures are distinct enough to be kept when assembling the final database. (**B**) Demonstration of the physical interpretation of principal axes in a 3D molecular structure. The principal axes x, y, and z are defined such that they each minimize the radial distribution of mass about successive orthogonal axes. The center image is a representation of the atomic positions from the structure on the left, with radii proportional to atomic masses. In this example, when viewed along the first principal axis (x, top right), there is very little radial distribution of masses about the central axis. In contrast, when viewed along the second principal axis (y, bottom right) the radial distribution of masses is in greater. The PMI are related to the magnitude of radial mass distribution about the respective principal axes, where increased radial mass distribution results in a higher moment.  (**C**) Mass-weighted radial atomic distance distribution for all 3D structures in the dmCCS database. Dashed lines indicate binning intervals used to compute binned radial mass distributions for individual structures as part of the MD3D features.

*2.8 – Table S1: Molecular Quantum Numbers (MQNs)*

| MQN | description | MQN | description |
|---|---|---|---|
| c | carbon atom count | hbdm | H-bond donor sites |
| f | fluorine atom count | hdb | H-bond donor atoms |
| cl | chlorine atom count | negc | negative charges |
| br | bromine atom count | posc | positive charges |
| i | iodine atom count | asv | acyclic monovalent nodes |
| s | sulfur atom count | adv | acyclic divalent nodes |
| p | phosphorus atom count | atv | acyclic trivalent nodes |
| an | acyclic nitrogen atom count | aqv | acyclic tetravalent nodes |
| cn | cyclic nitrogen atom count | cdv | cyclic divalent nodes |
| ao | acyclic oxygen atom count | ctv | cyclic trivalent nodes |
| co | cyclic oxygen atom count | cqv | cyclic tetravalent nodes |
| hac | heavy (non-hydrogen) atom count | r3 | 3-membered ring count |
| asb | acyclic single bonds | r4 | 4-membered ring count |
| adb | acyclic double bonds | r5 | 5-membered ring count |
| atb | acyclic triple bonds | r6 | 6-membered ring count |
| csb | cyclic single bonds | r7 | 7-membered ring count |
| cdb | cyclic double bonds | r8 | 8-membered ring count |
| ctb | cyclic triple bonds | r9 | 9-membered ring count |
| rbc | rotatable bond count | rg10 | ≥10-membered ring count |
| hbam | H-bond acceptor sites | afrc | nodes shared by ≥2 rings |
| hba | H-bond acceptor atoms | bfrc | edges shared by ≥2 rings |

*2.9 – Table S2: 3D Molecular Descriptors (MD3D)*

| **MD3D** | **description** |
|---|---|
| pmi1 | first principal moment of inertia |
| pmi2 | second principal moment of inertia |
| pmi3 | third principal moment of inertia |
| rmd02 | proportion of mass between 0 and 2 Å of center of mass |
| rmd24 | proportion of mass between 2 and 4 Å of center of mass |
| rmd46 | proportion of mass between 4 and 6 Å of center of mass |
| rmd68 | proportion of mass between 6 and 8 Å of center of mass |
| rmd8p | proportion of mass more than 8 Å from center of mass |
